# Versatile Stain Transfer in Histopathology Using a Unified Diffusion Framework

**DOI:** 10.1101/2024.11.23.624680

**Authors:** Xudong Yan, Mingze Yuan, Yao Lu, Ying Zhang, Zifan Chen, Peng Bao, Zehua Li, Bin Dong, Li Yang, Li Zhang, Fangxu Zhou

## Abstract

Histological staining is vital in clinical pathology for visualizing tissue structures. However, these techniques are laborious and time-consuming. Digital virtual staining offers a promising solution, but existing methods typically rely on Generative Adversarial Networks (GANs), which may suffer from artifacts and mode collapse. Motivated by the success of diffusion models, we present DUST, a novel **D**iffusion-based **U**nified framework for versatile **S**tain **T**ransfer in histopathology. To enhance domain awareness and task-specific performance, we propose a dual encoding strategy that integrates the stain types of both the source and target domains. Additionally, we introduce a dynamic dual-output head to address the unstable intensity issue encountered with conventional DDPM implementations. Validated on a curated four-stain kidney histopathological dataset (H&E, MT, PAS, and PASM), DUST demonstrates superior versatile stain transfer capabilities. Our research highlights the potential of diffusion models to advance virtual staining, paving the way for more efficient digital pathology analyses.

## 1. INTRODUCTION

Histological staining is crucial for visualizing tissue structures and cellular morphology in clinical pathology. Common stains like hematoxylin and eosin (H&E) provide essential contrast between nuclei and extracellular matrices. Special stains such as Masson’s trichrome (MT), Periodic Acid-Schiff (PAS), and Periodic Acid-silver Methenamine (PASM) are used to highlight collagen fibers, glycoproteins, and basement membranes, respectively. Despite their significance, these staining processes are time-consuming and labor-intensive, involving extensive sample preparation and manual execution by skilled histotechnicians. Moreover, given that the same section cannot be repeatedly stained, the necessity for multiple tissue sections for different stains poses challenges, especially in resource-constrained environments, hindering accessibility and increasing costs.

Recently, the advent of digital virtual staining has emerged as a transformative approach, enabling the conversion of one stain to another through deep learning [1]. Previous research has predominantly focused on developing specialized models to transfer H&E stains to one [2] or a few [1, 3] special stains, largely due to the prevalence of H&E staining. Yet, the concept of a unified framework that facilitates versatile stain conversion, *i*.*e*., translating any given stain to any desired stain (Figure 1 left), remains relatively unexplored. Developing such a framework is advantageous for two key reasons. First, despite H&E’s affordability, it has its limitations in providing detailed surface and color contrast, making the translation from special stains beneficial for capturing complementary details [1]. Second, the proven synergy of generalist models in both natural [4] and medical imaging [5] underscores the potential of a unified framework to surpass the capabilities of specialized models. By enabling efficient broad-spectrum stain conversion, this framework allows pathologists to instantly switch between different stain types within their existing workflows, thus supporting swift, effective, and high-throughput pathological analysis.

**Fig. 1.**
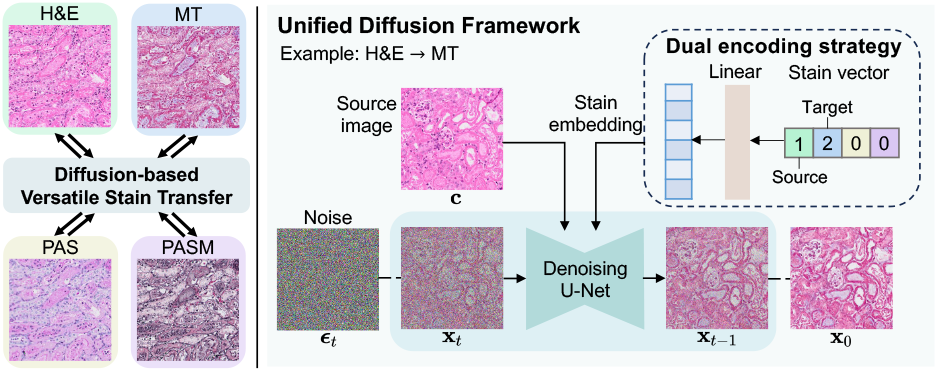
Framework Overview. Left: The versatile stain transfer problem, i.e., conversion across four stain domains: H&E, MT, PAS, and PASM. Right: Our proposed unified diffusion-based framework, which incorporates the source image and stain information as additional conditions. For stain conditioning, we introduce a dual encoding strategy that integrates information from both the source and target domains.

Furthermore, traditional stain transfer techniques [3, 6, 1, 7, 2] often rely on Generative Adversarial Networks (GANs) [8, 9, 10, 11], which can suffer from issues like artifacts and mode collapse. Additionally, unpaired translation methods [6, 3] lead to suboptimal results due to their inability to leverage pixel-wise correspondence, indispensable for detailed pathological images [7]. Fortunately, emerging diffusion models [12] have beat GANs in terms of stability and image quality across natural image generation [13, 14, 15], denoising, and translation [16] tasks, providing a promising alternative.

In this paper, we propose DUST, a novel **D**ffusion-based **U**nified framework designed for versatile **S**tain **T**ransfer in histopathology to address these limitations. Our method integrates the source image and the stain types of both the source and target domains as additional conditions to enable multitask learning. Unlike the commonly used one-hot encoding [10, 6, 3] or adaptive normalization strategies [17, 18], we propose a dual encoding strategy to integrate the stain types of both source and target domains, thereby enhancing its task-specific performance. Furthermore, we addressed the issue of unstable intensity encountered with conventional Denoising Diffusion Probabilistic Models (DDPM) implementations [12] by introducing a dynamic dual-output head [19]. Validated on our curated kidney histopathological four-stain (H&E, MT, PAS, and PASM) dataset, our method demonstrates superior performance on versatile stain transfer in the proof-of-concept transfer cycle. Our study underscores the potential of diffusion models in advancing the field of virtual staining.

## 2. METHOD

### 2.1. Unified Diffusion Framework

#### Problem Formulation

Versatile stain transfer can be formulated as a multi-domain image-to-image translation problem. Denote the set of histopathological stain domains with *K* stains as 𝒮 = *{*1, 2, …, *K }*. Given any source-target pair of stains *{s*_0_, *s*_1_}⊂ 𝒮 and a source image **x** with stain *s*_0_, we aim to apply a unified framework that can transfer the style of **x** from domain *s*_0_ to the desired domain *s*_1_, while preserving the content of **x**. The entire dataset is denoted as 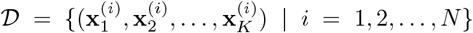, where 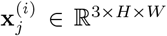 represents the *i*-th pathological slice with stain *j*. Here, we have *K* = 4 corresponding to the stains H&E, MT, PAS, and PASM.

#### Preliminaries

We briefly review key concepts of diffusion models [12]. Diffusion models initiate with a forward diffusion process that gradually adds Gaussian noise to an initial image **x**_0_, i.e., 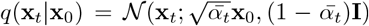, with 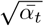 being hyperparameters. This process is reparameterized as 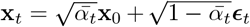, where ***ϵ***_*t*_ ∼ 𝒩 (0, **I**) represents the sampled noise. Then **x**_0_ can be easily backtraced via:

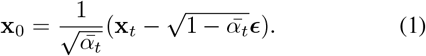

The reverse diffusion process, which reconstructs the original data by reversing the noise, is approximated by *p*_***θ***_(**x**_*t−*1_ |**x**_*t*_) = 𝒩 (**x**_*t−*1_; ***μ***_***θ***_ (**x**_*t*_, *t*), **Σ**_***θ***_(**x**_*t*_, *t*)). Training optimizes the variational lower bound on the log-likelihood of the initial data, simplified to ℒ _***θ***_ = *−* log *p*_***θ***_(**x**_0_|**x**_1_) + ℑ_*t*_ *𝒟 𝒟*_*KL*_ (*q*(**x**_*t−*1_ |**x** _*t*_, **x** _0_)*∥p*_***θ***_ (**x**_*t−*1_ |**x**_*t*_)). By characterizing the mean ***μ***_***θ***_ as a noise prediction network ***ϵ***_***θ***_ and fixing the variance [12], the training process simplifies to minimizing the mean-squared error between the predicted and actual noise, i.e.,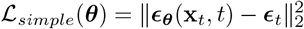.

#### Unified Diffusion Framework

To enable versatile stain transfer between any source-target stain pair, our framework accommodates additional inputs: the source image **c** from stain *s*_0_ ∈ 𝒮, the desired target stain *s*_1_ ∈ 𝒮 (where *s*_0_ *≠ s*_1_), the noise timestep *t*, and the noised target image 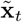. Given the sampled noise ***ϵ***_*t*_, our unified noise prediction network ***ϵ***_***θ***_ [12] denoises 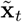 conditioned on these inputs, formalized by the optimization objective:

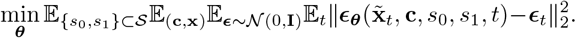

Here the source image **c** is conditioned via concatenation, and time features *t* are conditioned via sinusoidal positional encodings and spatial addition [15].

### 2.2. Advanced Functionality Design

#### Stain Conditioning

To achieve a unified framework, a crucial aspect is the conditioning on stain domains. We propose a dual-encoding approach to differentiate between various tasks. Specifically, we use an integer vector to denote both the source and target domains. We encode the source domain as 1 and the target domain as 2, i.e., the stain vector **r** = (*r*_1_, *r*_2_, …, *r*_*K*_), where 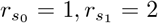, and *r*_*i*_ = 0 for each *i ∈ S \ { s*_0_, *s*_1_*}* (Figure 1 right). This vector is embedded and added with the time embedding to feed to the model through spatial addition, mirroring the class label conditioning approach in LDM [14]. This method incorporates both the source domain and the target domain into the model, enhancing its ability to distinguish between different tasks. The design can easily be extended to scenarios with multiple input domains.

#### Learning Objective

In a preliminary experiment, we observed that a conventional DDPM implementation [12] tends to produce unstable intensity shifts in the generated images, an issue evident in Figure 2(c). Closer inspection of the diffusion process revealed errors at larger timesteps *t*. This phenomenon is associated with the learning objective for noise prediction, particularly challenging at greater timesteps. Referring to Equation 1, when *t* is large, 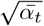 approaches zero, then even minor errors in ***ϵ*** can lead to substantial deviations in the final output, which causes the unstable intensity issue. To mitigate this issue, we incorporate the dynamic dual-output diffusion head [19] into our framework, which leverages the complementary strengths of noise prediction and original image reconstruction, resulting in more stable outputs (Figure 2(d)). Specifically, we modify the output channels of our model from *C* to 3 × *C*, where *C* represents the number of channels in the target images. For RGB images in our experiments, *C* = 3. The generate model **f**_***θ***_ computes:

**Fig. 2.**
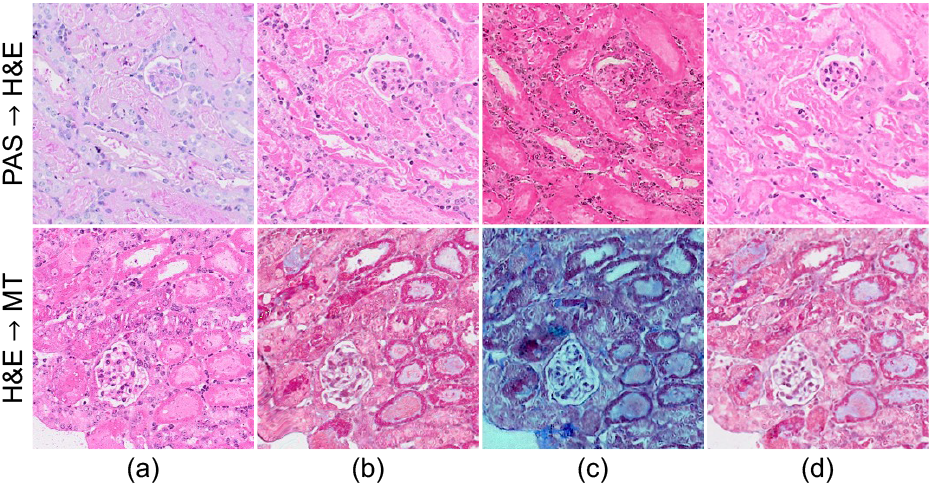
Image translation comparison: (a) Source, (b) Target, (c) DDPM [12], (d) Our method. The conventional DDPM implementation using a learning objective for noise prediction yields unstable intensity shifts, while our approach ensures stability.

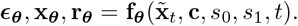

Here ***ϵ***_***θ***_ is the prediction of noise, **x**_***θ***_ is the prediction of the original image, and **r**_***θ***_ is a weighting factor. All of ***ϵ***_***θ***_, **x**_***θ***_, and **r**_***θ***_ have *C* channels. Subsequently, both **x**_***θ***_ and ***ϵ***_***θ***_ can estimate the forward process posterior mean 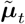. Specifically, ***μ***_**x**_(**x**_***θ***_) represents the mean obtained through the prediction of the original image **x**_***θ***_, while ***μ***_***ϵ***_(***ϵ***_***θ***_) represents the mean obtained through the prediction of noise ***ϵ***_***θ***_. We then use **r**_***θ***_ as a weight to combine these two estimates:

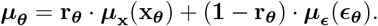

Finally, we define our loss function as:

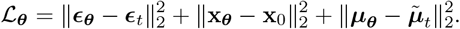

## 3. EXPERIMENTS

### 3.1. Experimental Settings

#### Datasets

We used Whole Slide Imaging (WSI) to obtain 40 slices of mouse kidney tissue for each staining method, including H&E, PAS, MT, and PASM. High-resolution scanned images with approximately ∼ 17,000 × 27,000 pixels were acquired using an Olympus VS200 microscope. The data were collected within the same batch, thus eliminating potential batch effects. To minimize pixel misalignment across various stains, we collected each eight slices in sequence and applied a four-stain cycle: H&E, PAS, MT, and PASM. Further, we employed VALIS [20], an advanced registration algorithm, for serial registration of multiple consecutive slices. The images were 4× downsampled and divided into 256 × 256 patches, following the approach described in [3]. Only patches containing a minimum of 20% foreground pixels were retained. After manual examination by expert pathologists, we created a pixel-level, well aligned paired dataset. For model training and evaluation, we randomly partitioned the dataset into two sets, with 29, 566 patches for training and 3, 532 for testing.

#### Implementation Details

We adopted a linear variance scheduler setting the maximum timestep *T* at 1, 000, with a noise level interval ranging from 0.002 to 0.02. Our model architecture is designed upon the efficient-UNet described in [13], with modifications tailored to our specific task. The architecture encompasses six stages, with channel numbers per stage being 32, 64, 96, 192, 256, and 512 respectively. Following [19], we adjusted the output head’s channel count from 3 to 9 to incorporate the dynamic dual-output head strategy. The dimensions for time and stain embedding were at 1024, incorporating a linear transformation for the stain vector embedding. We employed the AdamW optimizer with a constant learning rate of 1× 10^−4^. The training was conducted over 300 epochs for a unified model, accommodating all 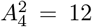 potential combinations of source-target domains. The batch size was 48. Our model was developed using PyTorch and MONAI frameworks and was trained on three NVIDIA A800 80G GPUs. For sampling, we used an advanced sampler, DPM-Solver++ [21], employing a single-step mode with 20 steps, translating a 256 × 256 image using ∼ 0.7s on a single A800 GPU.

#### Evaluation Metrics

Given the challenges in obtaining exact ground truth matches, even after registration [7], and the variability in stains produced by the same technician [1], we employed unpaired metrics in our experiments. Specifically, we utilized the Fréchet Inception Distance (FID) and the Kernel Inception Distance (KID). It is important to note that methods like SSIM, which evaluate structural similarity, are not well-suited for consecutive tissue slices. This is because consecutive slices inherently exhibit structural differences. Therefore, we opted for widely-used metrics in generative tasks, such as FID and KID, which are commonly accepted as standard evaluation metrics in the field. These metrics provide a robust assessment of the quality and realism of generated images, aligning with the current best practices in the community.

#### Baselines

Our comparative analysis includes a variety of baseline methods, encompassing both GAN-based strategies (such as pix2pix [8], CycleGAN [9], StarGAN [10], and StarGANv2 [11]) and a diffusion-based approach (Palette [16]). For methods not inherently equipped for multi-domain tasks (specifically pix2pix, CycleGAN, and Palette), we conducted individual training for each domain pair. Each model was developed using official source code and trained for 300 epochs.

### 3.2. Quantitative & Qualitative Results

#### Our Framework’s Proficiency in Versatile Stain Transfer

Utilizing a conceptual conversion cycle (H&E→ MT → PASM → PAS → H&E), we demonstrated our framework’s superiority in versatile stain transfer. As indicated in Table 1, our method surpasses competing techniques significantly in FID and KID metrics, recording an average FID of around 42 and a KID of approximately 0.02.

**Table 1.** Quantitative results on versatile stain transfer, using a proof-of-concept translation cycle: H&E → MT→ PASM → PAS → H&E. (FID: Fréchet Inception Distance, KID: Kernel Inception Distance)

|  | H&E→MT |  | MT→PASM |  | PASM→PAS |  | PAS→H&E |  |
| --- | --- | --- | --- | --- | --- | --- | --- | --- |
| Method | FID↓ | KID↓ | FID↓ | KID↓ | FID↓ | KID↓ | FID↓ | KID↓ |
| pix2pix [8] | 182.75 | 0.195 | 225.42 | 0.211 | 142.30 | 0.137 | 73.78 | 0.058 |
| CycleGAN [9] | 54.03 | 0.032 | 77.17 | 0.045 | 45.59 | 0.025 | 189.65 | 0.208 |
| Palette [16] | 87.35 | 0.058 | 51.88 | **0.012** | 75.20 | 0.064 | 78.10 | 0.052 |
| StarGAN [10] | 84.45 | 0.068 | 144.80 | 0.129 | 99.30 | 0.083 | 87.65 | 0.072 |
| StarGANv2 [11] | 104.51 | 0.099 | 151.68 | 0.190 | 87.71 | 0.083 | 89.30 | 0.079 |
| **Ours** | **41.76** | **0.019** | **42.66** | **0.012** | **41.92** | **0.022** | **41.46** | **0.023** |

Our method, in contrast to Palette [16], which is another diffusion-based approach that trains separate networks for each task, exhibits superior performance across all tasks. Notably, it outperforms Palette in the H&E to MT translation task, achieving an FID improvement of approximately 45.59. Additionally, our method demonstrates enhanced detail capture, possibly due to the introduced dynamic dual-output head. For instance, while Palette struggles with accurately representing cell nuclei and glomerulus boundaries, as observed within the red box in the fourth row of Figure 3(e), our framework achieves precise translations. This underscores the synergistic effect of our all-in-one pathological translation framework over task-specific models, resonating with the emerging trends in generalist and foundation models [5, 4, 22].

**Fig. 3.**
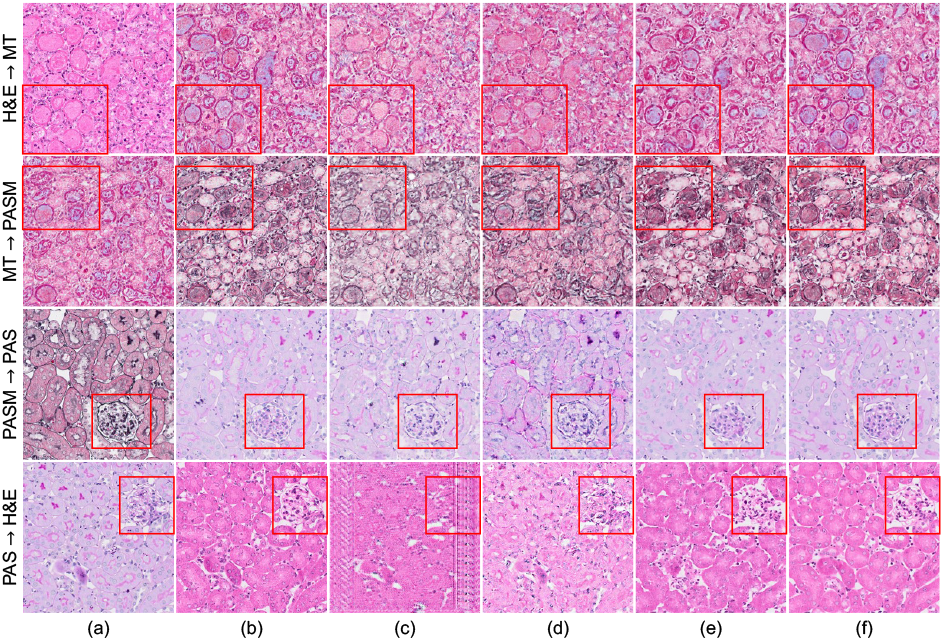
Visualization of image translation results using various methods in the proof-of-concept cycle: H&E → MT → PASM → PAS → H&E. From left to right: (a) Source, (b) Target, (c) CycleGAN [9], (d) StarGAN [10], (e) Palette [16], (f) Our method. Our approach effectively captures the target domain’s style while retaining the source image’s content. (Best viewed when zoomed in.)

In comparison to other multi-domain methods like Star-GAN [10] and StarGANv2 [11], our framework maintains consistent performance across all tasks, unlike the others which may falter in specific conversions, such as MT→ PASM, and struggle with style capture (see Figure 3(d)). This demonstrates that our method can maintain stability across multiple domains while using a unified framework.

### 3.3. Ablation Studies

We conducted an ablation study focusing on the design of stain conditioning. Besides our dual-encoding strategy, we employ a one-hot encoding scheme. In this method of stain conditioning, the source domain is disregarded, and a single number is used to represent the target domain *s*_1_. Our results demonstrate that dual-encoding yields superior performance compared to one-hot encoding. As detailed in Table 2, our “dual-encoding” strategy outperforms “one-hot encoding” strategy in all the four tasks. Our dual-encoding strategy, by integrating source stain information, enhances task differentiation within the framework. Consequently, we have selected the dual-encoding approach for our framework.

**Table 2.** Ablation studies comparing different strategies of stain conditioning.

|  | H&E→MT |  | MT→PASM |  | PASM→PAS |  | PAS→H&E |  |
| --- | --- | --- | --- | --- | --- | --- | --- | --- |
| Stain conditioning | FID↓ | KID↓ | FID↓ | KID↓ | FID↓ | KID↓ | FID↓ | KID↓ |
| 1) One-hot enc. | 42.28 | 0.020 | 47.86 | 0.020 | 52.19 | 0.035 | 49.05 | 0.033 |
| 2) **Dual enc.** | **41.76** | **0.019** | **42.66** | **0.012** | **41.92** | **0.022** | **41.46** | **0.023** |

## 4. CONCLUSION

We propose a novel diffusion-based unified framework for versatile stain transfer (DUST) in histopathology. By integrating dual encoding strategies and a dynamic dual-output head, it achieves superior performance in transferring images across multiple stain types. Our extensive experiments on a four-stain kidney histopathological dataset showcase its potential to advance the field of virtual staining towards efficient, accessible and high-throughput pathological analysis.

## 5. COMPLIANCE WITH ETHICAL STANDARDS

This study was performed in line with the principles of the Declaration of Helsinki. Approval was granted by the Ethics Committee of Peking University First Hospital (approval number: J2022138).

## Notes

### Competing Interest Statement

The authors have declared no competing interest.

